# Physical activity modulates early visual response and improves target detection in humans

**DOI:** 10.1101/2024.07.10.602924

**Authors:** Tom Weischner, Xinyun Che, Paul Schmid, Christoph Reichert, Annemarie Scholz, Robert T. Knight, Stefan Dürschmid

**Author notes:** Corresponding author: Stefan Dürschmid, Brennecke Str. 6, 39118 Magdeburg. **Author contributions:** S.D. conceived and designed the experiment. T.W. collected the MEG data. X.C., P.S., C.R., A.S., T.W., and S.D. analyzed the data, X.C., P.S., C.R., R.T.K., A.S., T.W., and S.D. interpreted the data. T.W., R.T. K., and S.D. wrote the manuscript. **Competing Interest Statement:** no competing interests.

## Abstract

Brain state changes affect visual perception by altering spatial resolution. Attention enhances the spatial resolution decorrelating neuronal activity in early nonhuman primate (NHP) visual cortex. Physical activity (PA) amplifies these attentional effects in rodents but impact of PA on visual perception in humans remains uncertain. We investigated the relationship between broadband high-frequency activity (BHA: 80-150 Hz) recorded with magnetoencephalography (MEG) and visual detection performance. We found that PA enhanced visual target detection predicted by a reduction of early BHA responses (<90 msec). This effect may be due to reduced interneuronal correlation to improve spatial resolution. Moreover, PA improved spatial integration time, as indicated by a linear relationship between reaction times and BHA variation with target eccentricity. These findings provide evidence that PA influences neuronal activity critical for early visual perception, optimizing visual processing at the initial stages of the visual hierarchy.

## Introduction

The perception of sensory stimuli is modulated by the behavioral state^1^ with attention improving stimulus perception by enhancing activity in relevant neuronal populations. This effect is accompanied by reduced interneuronal correlations with background neuronal populations^1–3^. This leads to enhanced spatial resolution and sharpens the focus of attention in nonhuman primates (NHP). In mice, physical activity (PA) leads to a brain state change that increases attention by boosting firing rates in V1 neurons ^4–7^, reducing the time needed for information accumulation in primary visual cortex^8^. How PA induced brain state changes impacts visual perception and early visual neuronal responses in humans remains uncertain. Previous studies in humans found increased activity across various neurophysiological components^9–11^ including SSVEP^11^ and alpha power modulation^10^. However, there is often a lack of behavioral improvement^12^ or even a decline in performance^13^ during movement, attributed to dual-task effects^11^ or head movements relative to visual stimulation^14^. Furthermore, research in humans has predominantly relied on EEG, focusing on low-frequency components, which decrease in contrast to increased neuronal firing with movement in rodents.

In humans, broadband high-frequency activity (BHA; 80 – 150 Hz), reflecting local neuronal population dynamics ^15,16^, is a comparable metric to V1 firing rate in rodents. Intracranial recordings show that the BHA tracks physical variation in simple grating stimuli with a rapid response modulation (<200 msec)^16^. In a visual detection experiment we asked subjects to detect a target defined by an orientation different from distractors. We hypothesized that differences in orientation can be better detected by increasing spatial resolution and sharpening attentional focus following PA. We aimed to determine whether BHA serves as an early indicator of the selection of targets embedded in distractors. Most importantly, we test whether PA amplifies this association. Given the prior NHP findings we hypothesized that PA would reduce interneuronal correlation^3^ and hence decrease the amplitude of BHA. We found that PA enhances target detection, coupled with a reduction in BHA (<90 msec). Importantly, this decline in BHA can be attributed to a selective decrease in target trials.

## Results

### Procedure

31 subjects participated in a target detection experiment after providing their written informed consent (see ***Figure 1A***). One subject was excluded due to excessive artifacts. Before each experimental block, participants were either instructed to rest or to use an MEG compatible pedal trainer for 2 min (see ***Figure 1A***). The pedal trainer permitted independent forward and backward movements (see ***Methods***). Participants were asked to move at a moderate speed, adjusting their pace individually, akin to a walking motion. Participants pseudo-randomly either started with the PA condition or the rest condition (counterbalanced across subjects). At the start of each trial, a fixation cross was displayed. Following a delay of 750 ms (± 250 ms), participants were presented with a visual search display below the fixation cross for 100ms. The target was defined by the degree of change in orientation angle from the distractor stimuli. In 50% of the trials of a block the target was presented at one of the 18 positions alongside 17 distractor stimuli (target trials). In the other 50% of trials only 18 distractors were shown (non-target trials). Target-and non-target trials were pseudo-randomly presented. Participants were instructed to indicate via button press whether a target was present (using their right middle finger) or absent (using their right index finger).

**Figure 1:**
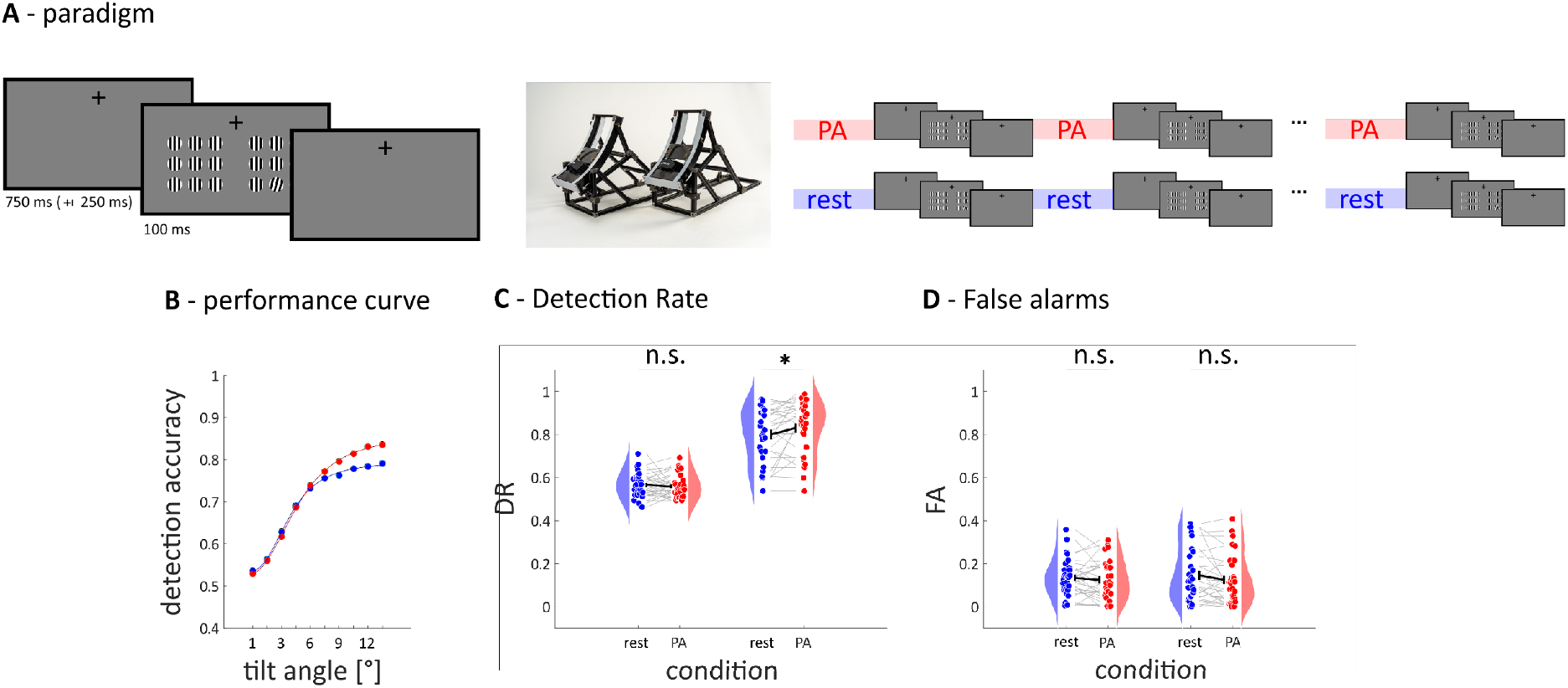
Depiction of the experimental paradigm and behavioral results. ***A*** in each trial, participants were asked to indicate whether a target stimulus defined by an angular difference relative to distractors was presented or not. Before each block, they rested or were engaged in movement. ***B*** shows the detection accuracy as a function of the angular difference. ***C*** shows that suprathreshold stimuli are better recognized when participants moved beforehand. ***D*** indicates a tendency of decreasing false alarms.

### I Behavioral results

First, we evaluated the target detection accuracy metrics detection rate (DR) and false alarm rates (FA). DR is defined as the probability of correctly reporting the target when it was presented for each block. In both conditions, we observed an increase in performance with the strength of the orientation angle from distractors (see ***Figure 1B***). We compared DR for subthreshold and suprathreshold targets (see ***Methods***). We averaged the three lowest (subthreshold; 1° - 4°, mean DR = 56.26 %; *SD* = 4.86 %) and the three largest orientation angles (suprathreshold; 11.5° - 14.5°, mean DR = 81.54 %; *SD* = 11.66 %; see ***Figure 1B***) separately for the rest and PA conditions. DR differed between conditions for suprathreshold targets (DR_rest_ = 80.04%; DR_PA_ = 83.05%; *t*_29_ = 2.2; *p* = .036) but not for subthreshold targets (DR_rest_ = 56.61%; DR_PA_ = 55.89%; *t*_29_ = .74; *p* = .464; see ***Figure 1C***). To ensure target detection improvement was not driven by an increased number of button presses, we compared FA (percentage of target responses when no target was present) for subthreshold and suprathreshold trials using a t-test. In contrast to DR, we found no difference in the false alarm rate neither for suprathreshold (FA_rest_ = 14.61%; FA_PA_ = 12.53%; *t*_29_ = 1.69; *p* = .103) nor subthreshold trials (FA_rest_ = 13.33%; FA_PA_ = 12.52%; *t*_29_ = .75; *p* = .453; see ***Figure 1D***). The trend in decreased false alarm rate after PA in the suprathreshold stimuli suggests that the participants reliably detected the targets rather than simply pressed the target button more frequently. Taken together, we found that PA improved visual target detection.

### II Broad Band High Frequency Activity

In the next step, we examined whether amplitude modulation of BHA predicts changes in performance. We band-pass filtered the time series in the high frequency range (80–150 Hz) and calculated the analytic BHA amplitude by Hilbert-transforming the filtered data for each trial and magnetometer. We found strongest BHA modulation in two magnetometers covering the occipital cortex bilaterally (z_max_ = 48.06 at 96 milliseconds; see ***Figure 2A***). We averaged BHA modulation across these channels and analyzed the effect of BHA amplitude modulation on performance and compared the BHA response in the PA with the rest condition.

**Figure 2:**
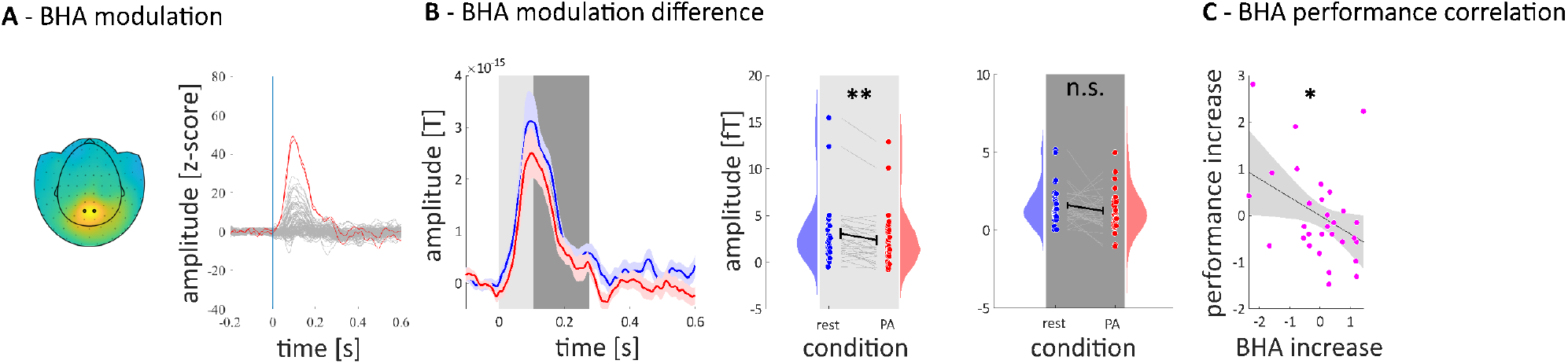
Depiction of broad band high-frequency activity (BHA) modulation. ***A*** shows the topographic distribution of BHA modulation. The two highlighted MEG sensors exhibit the strongest modulation after stimulus onset (red lines right). ***B*** demonstrates that PA leads to a reduction in the onset response of the BHA. ***C*** shows that the stronger the reduction in BHA during this rising phase, the better the performance of the participants.

### III BHA modulation following PA and rest

To analyze whether BHA differed between PA and rest, we categorized the BHA responses according to the two conditions and computed averages across all trials for each participant and condition. We examined the PA effect in both the increasing (0 to 96 msec) and the decreasing flank (96 to 250 msec; see ***Figure 2B***) of the BHA. We found a stronger response following rest compared to PA in the increasing (BHA_rest_ = 3.06 fT, BHA_PA_ =2.39 fT, *t*_29_ = 3.32, *p* = .0024) but not during the decreasing flank of the BHA (BHA_rest_ = 1.61 fT, BHA_PA_ = 1.24 fT, *t*_29_ = 1.62, *p* = .12; see ***Figure 2B***). This indicates that PA significantly decreases the amplitude of the early visual BHA response (<100 msec).

### IV BHA performance correlation

In the next step, we examined whether an amplitude decrease in the increasing flank of the BHA following PA explains the improved target detection. We determined both the difference in BHA and performance between rest and PA and found a significant correlation in the time domain of the increasing flank (*r*_max_ = -.39; *p* = .031; see ***Figure 2C***). This indicates that a reduction in BHA amplitude tracks performance improvement.

### V Distinction between target and non-target trials

Next, we examined whether participants exhibited different response patterns in target vs. non-target trials and whether these response patterns were influenced by PA. When we compared reaction times, we found a significant main effect of trialtype (F_1,28_ = 6.78; p = .0104; see ***Figure 3A***) due to longer reaction times in non-target (averaged across rest and PA: RT_NT_ = 457.0 msec) compared to target trials (RT_T_ = 404.7 msec). Using directed t-tests under the assumption that PA decreased reaction times we observed a reaction time decrease with PA both for target trials (RT_rest_ = 419.5 msec; RT_PA_ = 389.9 msec; t_29_ = 1.71; p = .049) and nontarget trials (RT_rest_ = 474.1 msec; RT_PA_ = 439.9 msec; t_29_ = 1.76; p = .044). BHA showed no difference between conditions in non-target trials (BHA_rest_ = 2.352 fT; BHA_PA_ = 2.210 fT; *t*_29_ = .45; *p* = .653) but a significantly reduced amplitude following PA compared to rest in target trials (BHA_rest_ = 2.656 fT; BHA_PA_ = 1.670 fT; *t*_29_ = 3.07; *p* = .0046; see ***Figure 3B***). Furthermore, only in target trials following PA, the BHA was correlated with RT (*r*_PA_target_ = .34, *p* = .067; *r*_rest_non-target_ = -.04, *p* = .799; see ***Figure 3C***) as mentioned above (*I – Behavioral results)*. In sum, PA reduces the BHA selectively in target present trials when interneuronal decorrelation is possible.

**Figure 3.**
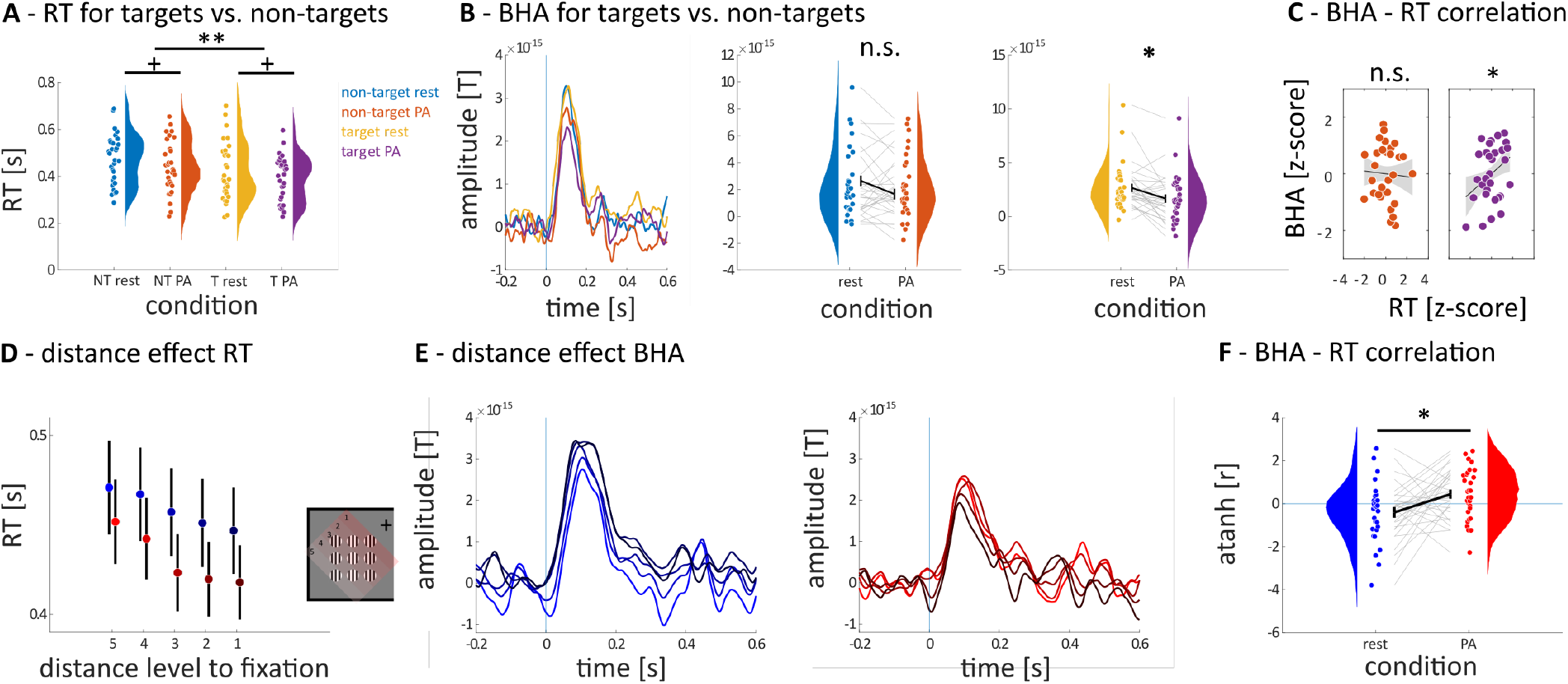
Depiction of BHA and RT patterns for different types of visual search. ***A*** shows that reaction times in target are shorter than in non-target trials. ***B*** shows that BHA on targets is reduced after PA compared to rest. ***C*** shows that the amplitude reduction after PA is associated with RT gain. The stronger the reduction, the shorter the RT. ***D*** shows that RT increases for targets further away from the fixation point. ***E*** shows differences in the modulation of BHA as a function of distance to fixation. ***F*** shows that after PA, the strength of BHA across different spatial distances correlates with RT for targets at different distances.

### VI Response modulation with distance to fixation

In the last step, we investigated whether PA impacts the spatial search efficiency by investigating whether the proximity of the target to fixation influenced reaction times. When comparing RT across different distance levels (see ***Figure 3D***), we observed a significant decrease in RTs in the PA condition for targets closer to the fixation point (summarized results in ***Table 1***) not evident in the rest condition. Next, we tested whether RT could be predicted by BHA amplitude (see ***Figure 3E***). We averaged RT and BHA response separately for each participant to targets presented at the five distance levels. We then correlated the resulting BHA values with the five average RT values separately for each subject. We then compared the resulting Pearson’s correlation values between conditions. First, reaction time – BHA correlation values were not different from zero in the rest condition (*r*_mean_ = -.19; *t*_29_ = 1.62; *p* = .12; see ***Figure 3F***). Conversely, in the PA condition, *r* values significantly differed from zero (*r*_mean_ = .24; *t*_29_ = 2.052; *p* = .049; see ***Figure 3F***) and were higher than in the rest condition (*t*_29_ = 2.45; *p* = .020). Taken together, we found that the spatial search efficiency as expressed by reaction time variations with eccentricity can be predicted by the BHA modulation following PA.

**Table 1:**
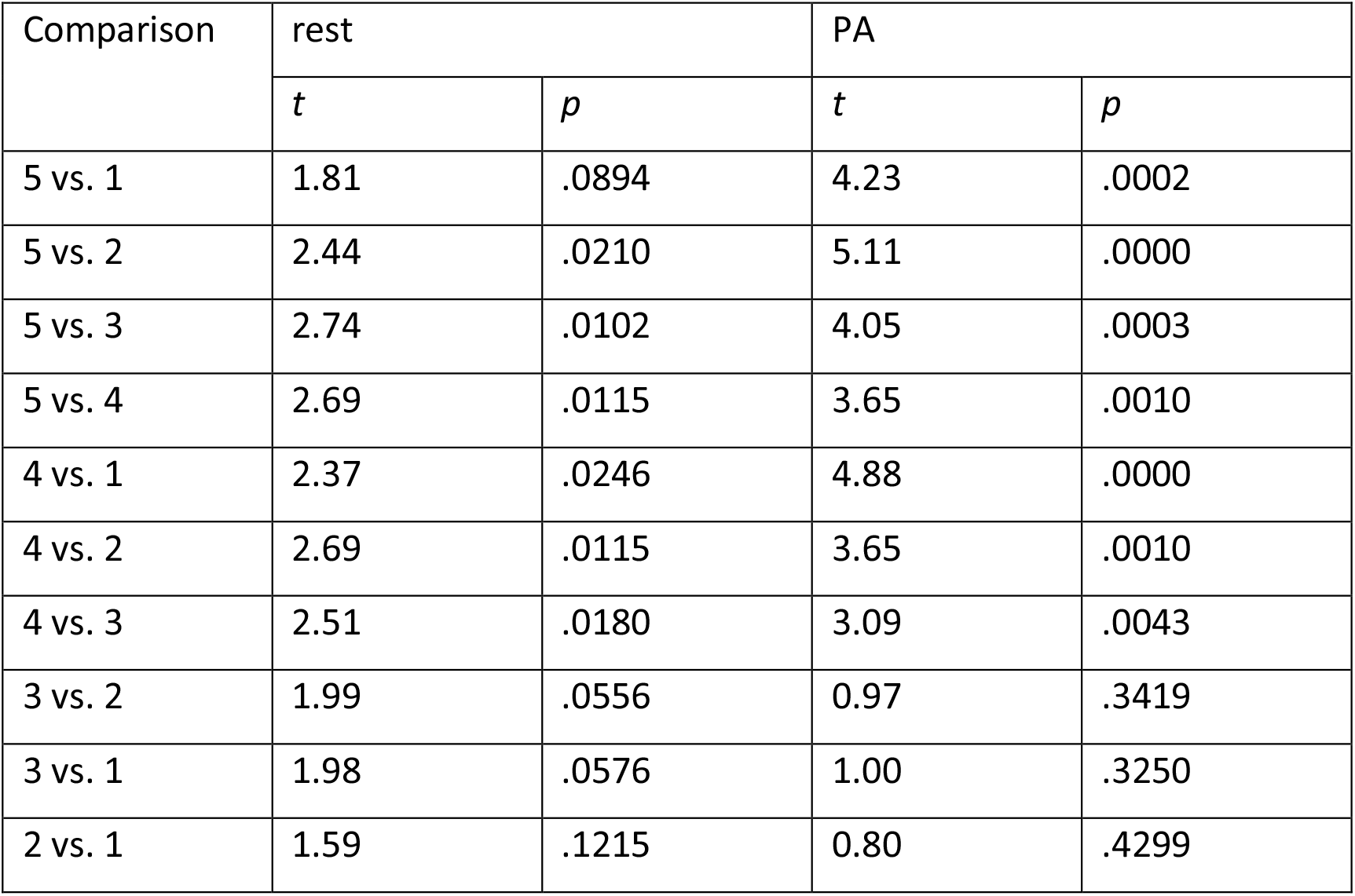
Results of RT probabilities in the proximity of the target to fixation. A decrease in RT in the PA condition for targets closer to the fixation point was observed, when comparing RT across different distance levels. However, this was not evident in the rest condition. Numbers 1 to 5 being the different distance levels (see ***Figure 3D*** for clarification).

| Comparison | rest |  | PA |  |
| --- | --- | --- | --- | --- |
|  | *t* | *p* | *t* | *p* |
| 5 vs. 1 | 1.81 | .0894 | 4.23 | .0002 |
| 5 vs. 2 | 2.44 | .0210 | 5.11 | .0000 |
| 5 vs. 3 | 2.74 | .0102 | 4.05 | .0003 |
| 5 vs. 4 | 2.69 | .0115 | 3.65 | .0010 |
| 4 vs. 1 | 2.37 | .0246 | 4.88 | .0000 |
| 4 vs. 2 | 2.69 | .0115 | 3.65 | .0010 |
| 4 vs. 3 | 2.51 | .0180 | 3.09 | .0043 |
| 3 vs. 2 | 1.99 | .0556 | 0.97 | .3419 |
| 3 vs. 1 | 1.98 | .0576 | 1.00 | .3250 |
| 2 vs. 1 | 1.59 | .1215 | 0.80 | .4299 |

## Discussion

We investigated whether physical activity impacts visual perception by improving spatial resolution and integration. Specifically, we examined whether physical exercise influences the behavioral detection of simple grating stimuli embedded in distractors. PA increased the target detection rate (DR) but did not alter the false alarm rate (FA). The early broadband high-frequency response (BHA) decreased in amplitude during successful target detection. PA improved spatial integration time, showing a linear gradient in the relationship between reaction times and BHA with distance to fixation. Our results are consistent with the more general view that sensory neuronal responses not only depend on stimulus strength, but also on behavioral state changes through physical exercise ^17–23^.

Research in mice reports improved perceptual performance during PA, yet human studies are inconclusive. Benjamin et al.^13^ explored how human perceptual performance changes during movement. Participants were asked in a series of contrast discrimination/detection tasks to identify the stimulus with higher contrast. Surprisingly, the authors discovered a contrary effect: thresholds in stationary conditions were lower compared to locomotion^13^. This discrepancy between human and rodent data could be explained by the experimental setup. Human subjects’ movement on the treadmill might induce vertical motion, causing a displacement in the visual field relative to stimuli optimally positioned only during the swing phase^14^. In our study, we eliminated the possibility of visual field shifts relative to the display since participants’ head movements are limited in the MEG. Furthermore, the task was carried out shortly after PA. Recent studies have examined whether walking or light exercise can improve visual processing compared to stationary conditions in humans. Various paradigms have been employed, such as spatial attention tasks^9^ and tasks eliciting SSVEP^11^ or alpha power modulation^10^. These investigations all observed a stronger modulation of the examined component with PA. However, none of these studies demonstrated a corresponding change in behavior. The lack of improvement could be because the employed tasks were relatively easy. However, some studies have shown a decline in performance^10,11,24^. This decline may result from a dual-task effect or from head movement relative to visual stimulation^14^. In summary, the inconsistent findings in human studies arise from the effects of movement in relation to visual stimulation and dual-task demands. To address these aspects, we separated movement from the cognitive task and demonstrated that PA improves visual perception.

Furthermore, we found an early effect (<100 ms) in the BHA. In contrast, previous studies in humans reported predominantly late neurophysiological effects (>200 ms^9,10^). This time frame does not align with the early V1 modulation observed in mouse research. In contrast to low-frequency activity in EEG, we examined BHA in MEG. MEG captures tangential activity, and high frequencies such as BHA exhibit minimal spatial spread. Our results demonstrate that PA improves detection of suprathreshold stimuli by modulation of activity early in the visual hierarchy.

Studies in mice demonstrated a doubling of visual neuronal response amplitudes without affecting firing rate or tuning properties^4^. However, findings in marmosets suggest that movement tends to decrease V1 responses^25^. The impact of PA on early visual activity in humans, reflected by a reduction in BHA, may parallel the effect observed in marmosets.

We also observed an impact of eccentricity on visual processing. Behaviorally, we found increased RT as the target was presented further away from the fixation point. Most importantly, RT variation with target distance to the fixation point positively correlated with BHA amplitude modulation to different levels of target eccentricity. Previous human studies did not establish a measure for spatial integration of information improvement by PA, because previous studies had all stimuli equidistant from the fixation point^9^. Additionally, studies with human participants often investigate spatial attentional shifts using N2pc in EEG, where the N2pc tends to decrease with eccentricity, despite being considered a target selection response^26^. This discrepancy is interpreted as due to the distribution of receptive fields in ventral areas, suggesting that N2pc amplitude decreases with target distance from the midline since fewer neurons are activated in the periphery compared to those near the midline. In contrast, in primates, an increase in firing rate with eccentricity can be observed, indicating that attention enhances spatial integration by increasing the cell’s summation area^27^. In agreement with this study, we can show that the early modulation of BHA reflects this process of spatial visual processing which is improved by PA.

In sum, we showed that PA positively influences visual perception by neurophysiological changes at an early stage of visual processing. The selective reduction of high-frequency responses when targets are detected post-exercise and the enhanced spatial integration time provide evidence for the impact of PA on neural processes underlying visual perception. In conclusion, our results are consistent with the view that sensory neuronal responses are dependent, not just on stimulus strength, but also on behavioral state across species.

## Methods

### Participants

After providing their written informed consent, a total of 31 subjects (19 female, range: 20-39 years, *M*: 25.19, *SD*: 4.36) participated in the experiment. All participants reported normal or corrected-to-normal vision, and none reported any history of neurological or psychiatric disease. All recordings took place at the Otto-von-Guericke University of Magdeburg and were approved by the local ethics committee (“Ethical Committee of the Otto-von-Guericke University Magdeburg”). Each participant was compensated with money or course credit.

### Paradigm

In this study, we investigate the impact of PA on visual target detection in a demanding task. Specifically, we compared detection of visual target stimuli defined by grating orientation following PA and rest. Participants were asked to detect a target stimulus that deviated from surrounding distractors in orientation. In each trial, 18 stimuli were simultaneously presented, grouped in two blocks of nine stimuli each (arranged in a 3 × 3 grid, see ***Figure 1A)***, which were positioned to the left and right of the fixation point. The stimulus presentation was implemented using Matlab R2018b (Mathworks, Natick, USA) and the Psychophysics Toolbox Version 3 extension (Brainard, 1997; Kleiner et al., 2007). All stimuli were presented on a gray background and projected onto a semi-transparent screen with an DLP LED projector (PROPixx VPX-PRO_5050B, VPixx Technologies Inc., Saint-Bruno, QC, Canada), using a resolution of 1.920×1.080 pixels and a refresh rate of 60 Hz. Distance to the screen, on which the projection extended, was kept constant at 100 cm.

The movement in the PA condition of the experiment was achieved through a custom-made, nonmagnetic pedal-trainer, which was mounted to the dedicated chair in the MEG chamber. This trainer allows for the adjustment of both angle and height, customizing it to fit individual participant needs. The breaks between the blocks in which the pedal-trainer was to be used and the breaks in which the participants should only rest lasted about 120 s each. The visual detection task was to detect the presence of a target. The target stimulus was randomly visible in 50% of the trials. The distractors were uniformly vertically aligned, while the target’s angle varied between 1° and 14.5° across different blocks, with 10 steps of 1.5°. Each stimulus consisted of four parallel black stripes on a white background. The visible portion of each grating occupied 1° 51’ of visual angle. One whole 3×3 grid of gratings occupied approximately 7° of visual angle, with 44’ between the gratings. The space between the two grids was measured at 3° 22’, the space between the lower end of the fixation point and the upper end of the gratings at 1° 25’. Whereas the fixation point itself had a visual angle of 30’.

### MEG and Eye Movement Recordings

Participants were equipped with metal-free clothing and seated in a dimmed, magnetically shielded recording booth. Responses were given with the right hand via an MEG compatible VPixx response system (VPixx Technologies Inc., Saint-Bruno, QC, Canada). Acquisition of MEG data was performed with the participant in a sitting position, using a whole-head Elekta Neuromag TRIUX MEG system (Elekta Oy, Helsinki, Finland), containing 102 magnetometers and 204 planar gradiometers. Sampling rate was set to 1000 Hz. The vertical electrooculogram (EOG) was recorded using one surface electrode above and one below the right eye. For horizontal EOG, one electrode each on the left and right outer canthus were used. Additionally, participants’ eye movements were recorded using an Eyelink 1000 Plus system (SR Research) with a Long-Range Mount configuration and a built-in camera at a sampling rate of 500 Hz. We attached the Long-Range Mount configuration to the presentation screen, so that all participants were seated in the same distance to the eye tracking camera (100 cm). Prior to data collection, participants completed an eye tracker calibration with the built-in 9-point grid method. Preparation and measurement took about 2.5 h.

### Preprocessing and Artifact Rejection

Maxfilter was applied to remove external noise from the MEG recordings. MEG and EOG data were down sampled to 500 Hz before further preprocessing steps were applied. We used Matlab (Mathworks, Natick, USA) for all offline data processing. We involved only the 102 magnetometers in our analyses. All filtering (see below) was done using zero phaseshift IIR filters (fourth order; filtfilt.m in Matlab). First, we bandpass-filtered the data between 1 and 200 Hz. Then, we notchfiltered the data to discard line noise (50Hz and its 2nd, 3rd, and 4th harmonic). Data were epoched from 1s preceding the stimulus presentation to 2 s after the presentation onset. Each trial was then baseline-corrected relative to the 500 ms interval preceding the stimulus presentation. We then visually inspected all data, excluded epochs exhibiting excessive muscle activity, as well as time intervals containing artifactual signal distortions, such as signal steps or pulses. Trials exceeding a variance criterion of 4 times the mean variance were excluded, as well as trials with a reaction time < 0.1 s or > 1.5 s.

### Statistical Analysis

In general, we conducted the following analysis steps. First, we compared behavioral measures between PA and rest (*I – Behavioral Analysis*). This included target DR, FA and global reaction times.

Next, we analyzed the grand average MEG-BHA response (*II – Broad Band High Frequency Activity*). In the following step we compared BHA modulation between the two movement conditions (*III – BHA modulation following PA and rest*). In the next step, we examined whether amplitude modulation of BHA predicts changes in performance (*IV – BHA performance correlation*). We then explored whether participants exhibited different responses patterns depending on whether a selection is possible and whether these response patterns were influenced by PA (*V – Distinction between target and non-target trials*). In the last step, we investigated whether PA has an impact on the spatial search efficiency (*VI – Response modulation with distance to fixation*)

### I Behavioral Analysis

#### Target Detection

We determined the probability of correctly detecting the target in each block. We averaged the resulting detection rate across subjects both for the rest and PA condition and fitted a psychometric function

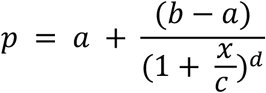

to the resulting values to model the relationship between stimulus variability (*x* - target orientation strength) and perceptual accuracy *p*. We defined the visual threshold across all participants as the stimulus values at which they responded correctly with 75% accuracy. As a first step, we compared the target detection rate (DR - % of correctly reporting the target when it was presented) for subthreshold (accuracy<75%) and suprathreshold targets (accuracy>75%). To have an equal number of trials, we averaged the three lowest (1° - 4°) and the three largest orientation angles (11.5° - 14.5°; see ***Figure 1B***) separately for the rest and PA conditions. We then compared resulting detection rates both for subthreshold and suprathreshold trials using a t-test.

#### False Alarm Rate

To rule out the possibility that an improvement in the target DR only occurred because the participants pressed the target button more frequently, we compared the FA (percentage of target responses when no target was presented) both for subthreshold and suprathreshold trials using a *t* test.

#### Reaction times

We grouped RT for trials with subthreshold and suprathreshold target angles and compared them using a *t* test between PA and rest.

### II Broad Band High Frequency Activity

For each trial and magnetometer, we band-pass filtered the time series in the broadband high frequency range (80–150 Hz) and obtained the analytic amplitude (called BHA) of the signal by Hilbert-transforming the filtered time series. We smoothed the resulting time series such that the amplitude at a given time point *t* was always the mean activity of 20 ms around this time point. The data were then baseline corrected by subtracting for each trial and each channel the mean activity of the time interval 100 ms preceding stimulus onset from every sampling point for each trial and each channel. Afterwards we identified the best two channels showing the most significant amplitude modulation in the BHA following the stimulus presentation. To that end, we normalized BHA amplitude values for each channel by dividing the BHA at each time point by the standard deviation. The two channels with highest z-scores in the interval ranging from 0 to 300 ms were labeled as stimulus responsive. For subsequent analyses we averaged the BHA data over these two MEG sensors (see ***Figure 2A***).

### III BHA modulation following PA and rest

In the next step, we examined whether the BHA differed between the PA and rest condition. We grouped the BHA responses based on the two conditions and averaged across all trials for each participant. Subsequently, we conducted *t* tests to compare the BHA separately during the increasing flank (see ***Figure 2A***) and the decreasing flank interval between the PA and rest conditions across subjects.

### IV BHA performance correlation

In the next step, we examined whether amplitude modulation of BHA predicted changes in performance. Generally, we determined both the difference in BHA between the two conditions and the performance difference between the two conditions. Specifically, we subtracted BHA_PA_ from BHA_rest_ at each time point for each subject, resulting in a ΔBHA time series for each participant. Furthermore, we subtracted DR_PA_ from DR_rest_, yielding a ΔDR value for each subject. These ΔDR values were then correlated with ΔBHA values averaged across the temporal interval of the increasing BHA. To correct statistical significance for multiple comparisons we applied Bonferroni correction. Since activity at each time point *t* depends on activity at time point *t*-1, two adjacent tests cannot be regarded as independent. Hence, we determined how many individual components are contained in the grand average BHA and corrected the alpha value by the number of components that significantly explained variance. We carried out a principal component analysis (PCA) and determined the eigenvalues of the resulting components. Components with an eigenvalue larger than 1 were considered to explain a significant amount of variance within our data. In the BHA activity we found 4 individual components. Hence, the corrected *P* value for the BHA is 0.05/5 = 0.01.

### V Distinction between target and non-target trials

Next, we examined whether participants exhibited different response patterns in target vs. non-target trials and whether these response patterns were influenced by PA. Initially, we compared RT to targets and non-targets after periods of rest or PA using a two-way analysis of variance (ANOVA) with trial type (target vs. non-target) and condition type (PA vs. rest) as factors. To accomplish this, we categorized trials based on trial type, averaged RT across trials for each participant, and separated them for each condition type. Subsequently, we explored whether PA had an impact on BHA in response to targets or non-targets. We conducted individual t-tests to compare BHA between PA and rest during the increasing flank interval. (see ***Figure 2A***) between the PA and rest conditions across subjects. Finally, we tested whether individual differences in BHA explained differences in RT. For this, we correlated the BHA averaged across the increasing flank interval with the individual RT separately for each trial type in each movement condition.

### VI Response modulation with distance to fixation

In the last step, we investigated whether target proximity to the fixation point influenced reaction times. Target positions were categorized into five distance levels, and corresponding trials were grouped accordingly. Average RT for each participant at these levels were compared using Bonferroni-corrected pairwise *t* tests. Following this, we explored the impact of target distance on BHA by organizing trials based on target presence for both rest and PA condition. We then correlated BHA amplitude with the five distance levels and individual RT. Resulting Pearson’s correlation coefficients were transformed (by calculating the inverse hyperbolic tangent) and compared between rest and PA conditions.

## Data availability

All data shown in results and figures can be accessed through https://github.com/SDuerschm/PEHFA

